# Stable isotope probing of carbonyl sulfide and cyanate pathways in microbial thiocyanate biodegradation

**DOI:** 10.1101/2024.12.18.629290

**Authors:** Yu-Chen Ling, Mathew P. Watts, John W. Moreau

## Abstract

Thiocyanate (SCN^−^) is toxic to many aquatic organisms at elevated concentrations. Historically, large amounts of SCN^−^ were released to the environment by gold mining and coal coking. Microbial SCN^−^ degradation provides a cost-effective approach to remediation; however, knowledge is still limited about the relative roles of the two known enzymatic SCN^−^ biodegradation pathways. Here we applied stable carbon and nitrogen isotope probing of SCN^−^-degrading microbial cultures to assess quantitatively the relative contributions of the cyanate (OCN^−^) and carbonyl sulfide (COS) pathways over a 24-hour experiment. In contrast to common assumptions, our results demonstrate that the OCN^−^ pathway initiates *before* the COS pathway and can contribute up to 70% of early SCN^−^ biodegradation. These results yield new insights into the dynamics of SCN^−^ biodegradation, and also show that microorganisms can incorporate carbon and nitrogen from the SCN^−^ into biomass, resolving the question of nitrogen mass balance. Our study holds implications for improving bioreactor design for SCN^−^ bioremediation and for understanding the dynamics for sulfur, carbon and nitrogen flows from SCN^−^ in natural environments into their respective biogeochemical cycles.

**Importance:** While the process of SCN^−^ biodegradation has been well studied, relative contributions from the two known enzymatic pathways are still poorly resolved, hindering our ability to optimize bioremediation systems treating SCN^−^-contaminated wastewater. Also, as SCN^−^ biodegradation in nature impacts the biogeochemical cycling of sulfur (as well as carbon and nitrogen), a more quantitative understanding of enzymatic SCN^−^ biodegradation pathways will yield insights into how the reactivity of this environmentally ubiquitous compound influences marine, terrestrial and atmospheric sulfur chemistry. Here we used carbon and nitrogen stable isotope probing to assess contributions to SCN^−^ biodegradation from known enzymatic pathways in a microbial consortium grown from SCN^−^-contaminated mine tailings. We found that the cyanate pathway dominates SCN^−^ biodegradation initially, before activation of the carbonyl sulfide pathway. We attribute this ordering to the greater amount of bioenergy conservable via the cyanate pathway during the earliest stages of bioremediation.

## Introduction

Thiocyanate (SCN^−^) is formed in many natural terrestrial and marine environments as a reaction product of aqueous bisulfide (HS^−^) with cyanide (CN^−^) produced by bacteria, algae and plants (1). Thiocyanate is also a product of reactions between CN^−^ and sulfide-containing minerals where CN^−^ solutions are commonly used to leach gold from ores (2–4), and is also a common pollutant from coal-coking processes (5–7). Indeed industrial sources have produced wastewater SCN^−^ concentrations as high as 2-3 g/L (8, 9), which is highly toxic to aquatic organisms (10–12), although microbial degradation of SCN^−^ also occurs naturally (13–15).

Microbial SCN^−^ biodegradation has been applied to mine waste remediation (16, 17). Previous studies report that mixed-species microbial communities degrade SCN^−^ more efficiently than pure cultures (18, 19). Recent work has illustrated experimentally that microbial consortia sourced from gold mine tailings can effectively bioremediate SCN^−^ contaminated mine waste autotrophically with only the addition of PO_4_^3-^ (20), in contrast to better-known but arguably less simplified industrial-scale and proprietary heterotrophic treatments (e.g., ASTER, (21)).

Microbial SCN^−^ degrading consortia have been shown to adapt to environmental stresses such as exposure to heavy metals (22), fluctuating temperatures (23, 24) and short-term changes in pH (25, 26). Their ability to withstand such excursions from ideal growth conditions makes them a robust tool for bioremediation.

Microorganisms degrade SCN^−^ via either of two known enzymatically-mediated processes, the carbonyl sulfide (COS) pathway (eqs. 1 and 2) or the cyanate (OCN^−^) pathway (eq. 3, (27)):

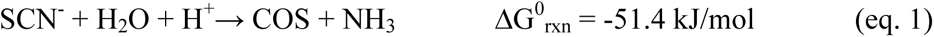

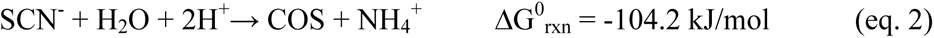

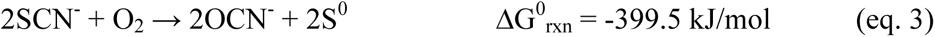

Since both ammonia (NH_3_) and ammonia ion (NH_4_^+^) are present at pH 7.5 in aqueous solutions under room temperature (28), we have listed both reactions (eqns. 1 and 2). The energy yield from the reactions (ΔG_rxn_) is calculated using the equation:

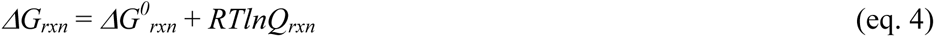

where *ΔG^0^_rxn_*is the standard free energy of the reaction, *R* is the universal gas constant, *T* is the temperature in Kelvin, and Q_rxn_ is the reaction quotient.

The standard free energy of reaction for each of these initial degradation steps is greater for the cyanate pathway. The products of both pathways, COS and OCN^−^, serve as energy sources for microorganisms, leading to the generation of H_2_S and NH_3_/NH_4_^+^:

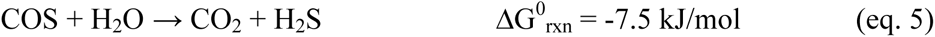

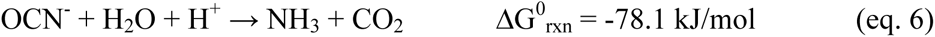

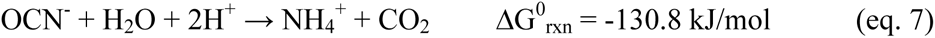

Interestingly, here the standard free energy for the cyanate pathway is still larger. However, subsequent oxidations of *all* reaction products (eqns. 1-3 and 5-7) exhibit the largest standard free energy for elemental sulfur (S^0^) oxidation (–1064 kJ/mol), followed by the oxidation of bisulfide (HS^−^, after deprotonation of H_2_S at most environmental pH values), ammonia (NH_3_) and ammonia ion (NH_4_^+^), yielding –749.6, –354.5 and –301.7 kJ/mol, respectively). Although calculation of the overall reaction energy yield must be refined based on chemical concentrations, the substantial standard free energy yield favoring *all* reaction products’ oxidation effectively “pulls through” SCN^−^ biodegradation via both the cyanate and carbonyl sulfide pathways. Sulfur-oxidizing bacteria then oxidize S^0^ and H_2_S to sulfate (SO_4_^2-^) under oxic conditions (23, 29, 30), while NH_4_ ^+^ may be an end product (23) or oxidized via nitrification during a two-stage treatment (17, 29). On the other hand, COS has lower solubility than OCN^−^ (0.125 g and 11.6 g in 100 mL water at 25°C, respectively), and only slowly degrades to H_2_S under ambient environmental conditions, which explains why most COS is volatilized (31). Volatilized COS contributes to the atmosphere as an important trace gas of sulfur with albedo-enhancing properties (32). However, there is limited knowledge regarding COS cycling in the environment, compared to other end products of SCN^−^ biodegradation.

Therefore, analyzing the dynamics of the COS and OCN^−^ enzymatic pathways for SCN^−^ degradation will inform the efficiency and sustainability of bioremediation systems as well as help us understand the role of microbially produced COS in the biogeochemical sulfur cycle. Furthermore, previous studies have noted a mass imbalance for nitrogen during SCN^−^ biodegradation, possibly due to incorporation of N from OCN^−^ into microbial biomass. Tracing nitrogen flow during SCN^−^ consumption would test this hypothesis. However, limited knowledge exists about the relative roles of, or potential for interplay between, the COS and OCN^−^ pathways for SCN^−^ biodegradation.

In this study, we used a microbial consortium enriched from a gold mine tailings storage facility (Fig. 1A) for laboratory experiments to investigate microbial SCN^−^ bioremediation (Fig. 1B). We tracked chemical changes to follow the biodegradation process and applied stable-isotope probing (SIP) to identify microorganisms involved in the COS pathway during SCN^−^ bioremediation (Fig. 1C). By adding unlabeled KSCN and labelled KS^13^C^15^N substrates, we isolated and sequenced both isotopically labelled (heavier) and unlabeled (lighter) DNA. The functional gene encoding for the COS pathway, *scnC* (33) was quantified using quantitative polymerase chain reaction (qPCR) (Fig. 1D). For the OCN^−^ pathway, researchers only relatively recently discovered the TcDH enzyme that mediates SCN^−^ biodegradation via intermediate OCN^−^ formation (34), and design of suitable primers to detect TcHD encoding gene is ongoing (35). Therefore, a proxy for OCN^−^ pathway activity was quantified by measuring OCN^−^ concentrations during SCN^−^ biodegradation (Fig. 1E). Finally, the amount of N incorporated into microbial biomass was measured and calculated using elemental analysis – isotope ratio mass spectrometry (EA-IRMS, Fig. 1F). Our results allow assessment of the relative performance of the two known pathways for SCN^−^ biodegradation in a mixed microbial community grown from mine waste.

**Figure 1.**
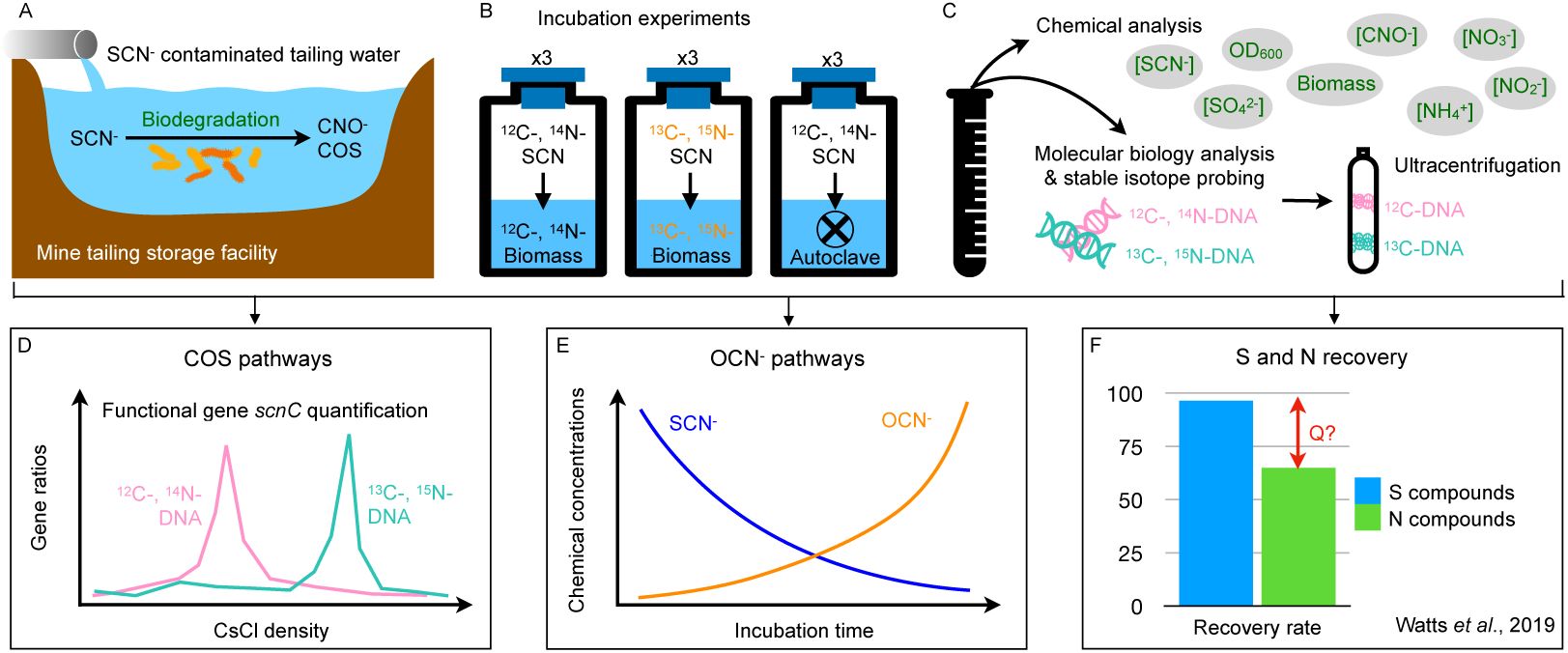
Schematic diagram of the workflow in this study. (A) The mine tailings storage facility contains thiocyanate (SCN^−^) contaminants, which microorganisms can degrade into cyanate (OCN^−^) and carbonyl sulfide (COS). (B) Microbial inocula were collected from the mine tailing storage facility, and triplicate enrichment experiments were conducted using SCN^−^ labeled with isotopes ^13^C and ^15^N, as well as non-labeled SCN^−^, to monitor the SCN^−^ biodegradation process. (C) Incubation sampling was designed for both chemical and molecular biological analyses. We aimed to (D) elucidate the SCN^−^ biodegradation process through the COS pathway by using quantitative polymerase chain reaction (qPCR) of *scnC* functional genes in isotope-labeled and non-labeled DNA, (E) investigate the SCN^−^ biodegradation process through the OCN^−^ pathway by comparing variations in SCN^−^ and OCN^−^ concentrations during SCN^−^ biodegradation experiments, and (F) test the hypothesis from (23) regarding the incomplete recovery of nitrogen during bioreactor culturing of SCN^−^ degraders.

## Results

### Chemical analyses from SCN-biodegradation experiments

Within 24 hours experimental time, SCN^−^ concentrations degraded from 5.5 ± 0.154 mM to below the detection limit (Fig. 2A), coupled with an increase in OD_600_ from 0.439 ± 0.003 to 0.493 ± 0.008 for experimental replicates (Fig. 2E). Compared with control groups where SCN^−^ was not degraded across the incubation period (Fig. 2A), the results confirmed that microorganisms mediated SCN^−^ degradation.

**Figure 2.**
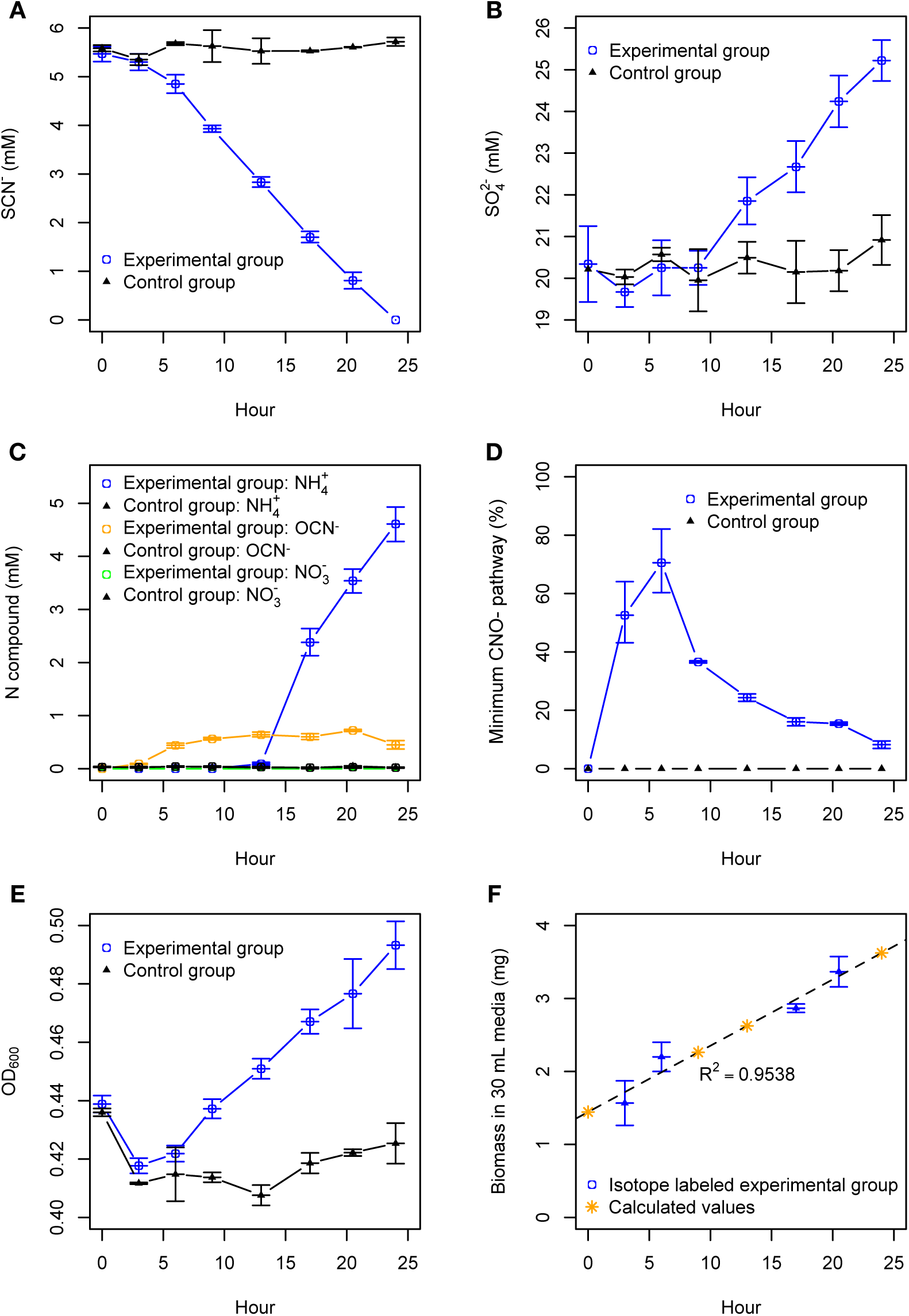
Chemical compositions of. (A) SCN^−^, (B) SO_4_^2-^, (C) N-compounds including NH_4_^+^, OCN^−^ and NO_3_^−^, (D) calculated contribution of OCN^−^ pathways, (E) OD_600_, and (F) measured and calculated biomass weight during the SCN^−^ degrading incubation experiments over time. Error bars present the standard deviation of mean values.

During the experiment, concentrations of OCN^−^ first increased at 3 hours (Fig. 2C), followed by an increase in SO_4_^2-^ and NH_4_^+^ concentrations after 13 hours (Figs. 2B and 2C). Concentrations of OCN^−^ were below the detection limit at 0 hours, increased to 0.09 ± 0.009 mM at 3 hours, 0.44 ± 0.041 mM at 6 hours, and 0.56 ± 0.025 mM at 9 hours, to reach a maximum range between 0.64 ± 0.046 and 0.72 ± 0.022 mM during 13-20.5 hours before decreasing after 24 hours (Fig. 2C). Concentrations of SO_4_^2-^ were between 19.67 ± 0.362 and 20.34 ± 0.908 mM from 0-9 hours, then increased to 21.85 ± 0.566 at 13 hours, 22.67 ± 0.619 at 17 hours, 24.24 ± 0.619 at 20.5 hours, and 25.22 ± 0.491 at 24 hours (Fig. 2B). Concentrations of NH_4_^+^ were below the detection limit from 0-9 hours, and then increased to 0.09 ± 0.026 at 13 hours, 2.38 ± 0.257 at 17 hours, 3.54 ± 0.224 at 20.5 hours, and 4.61 ± 0.324 at 24 hours (Fig. 2C). Concentrations of NO_3_^−^ were at background levels throughout the experiment (<0.05 mM).

Incorporation of ^15^N into microbial biomass was detected within 3 hours of initial incubations and exceeded 6 At%^15^N by 6 hours (Table 1). The weight of the biomass in 30 mL media increased from 1.6 mg at 3 hours to 2.2 mg at 6 hours, 2.9 mg at 17 hours, and 3.4 mg at 20.5 hours (Table 1). Measured biomass weights were fitted to a linear regression line with an R^2^ value of 0.95 (Fig. 2F). Based on this analysis, we calculated the theoretical biomass weights in 30 mL media were 1.4, 2.3, 2.3, and 3.6 mg at 0, 9, 13, and 24 hours, respectively.

**Table 1.**
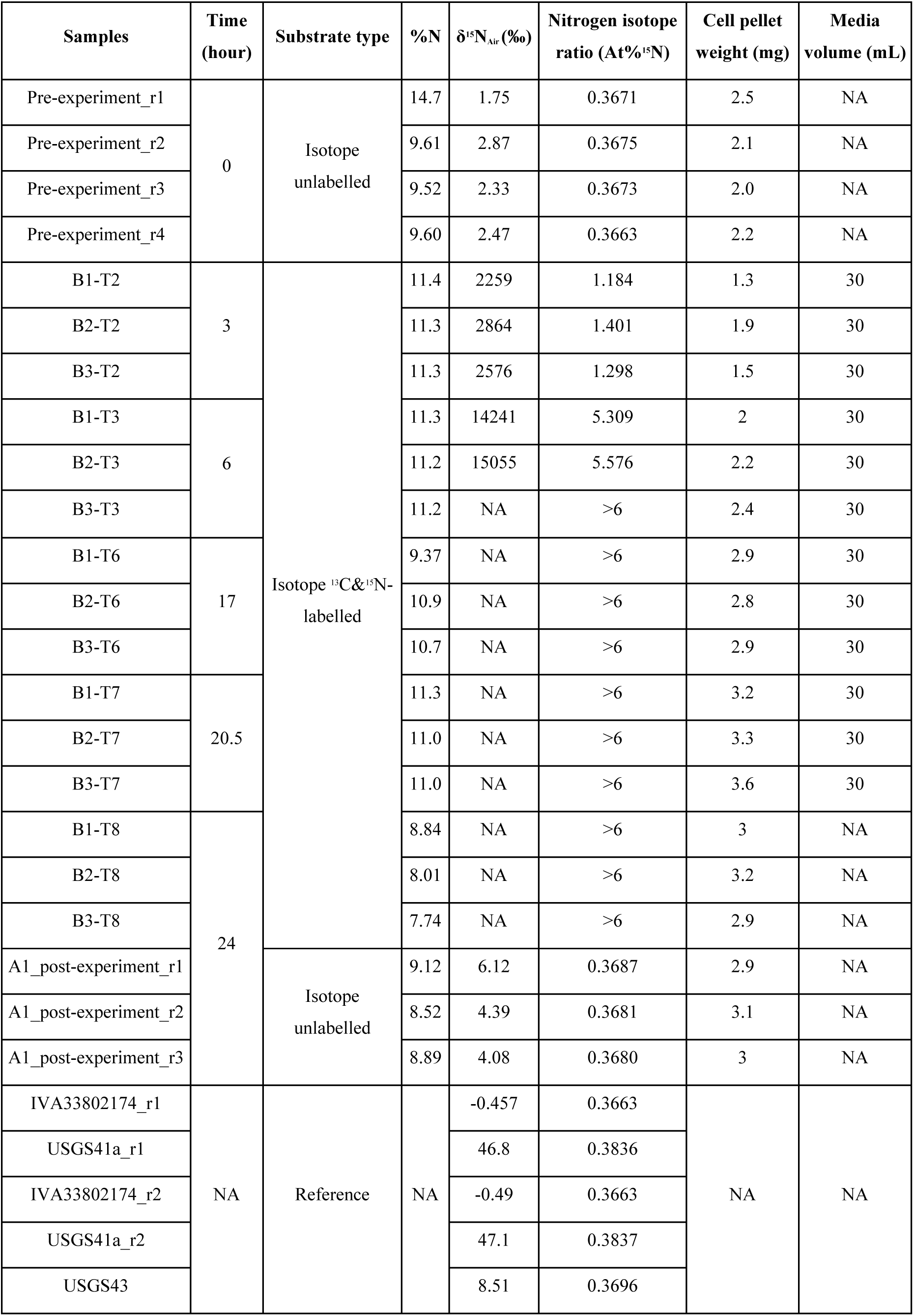
EA-IRMS measurement of total N and δ^15^N in microbial biomass during SCN^−^ degrading incubation experiments at T2, T3, T6, T7, before and after experiments.

### OCN^−^ pathway contribution

The OCN^−^ pathway contributions were calculated based on the amount of SCN^−^ converted into OCN^−^ at each time point. The OCN^−^ pathway made a higher contribution to SCN^−^ degradation at initial time points (T2 and T3), and the maximum contribution was up to 70.6% at 6 hours (Fig. 2D). Calculated OCN^−^ pathway contributions at other time points were 52.6% at 3 hours, 36.6% at 9 hours, 24.4% at 14 hours, 16.1% at 17 hours, 15.4% at 20.5 hours, and 8.2% at 24 hours. Since OCN^−^ can be decomposed through biotic and abiotic reactions (36), OCN^−^ pathway contribution rates calculated based on OCN^−^ concentrations would constitute minimum values and we have ignored the potential effect of OCN^−^ removal mechanisms here. Although calculated contributions from the OCN^−^ pathway decreased after T3 (6 hours, Fig. 2D), concentrations of NH_4_^+^ increased after T4 (9 hours, Fig. 2C). We are unable to distinguish, however, if NH_4_^+^ was generated through OCN^−^ degradation or via the COS pathway. Therefore, actual contributions from the OCN^−^ pathway could have been higher than the calculated values.

OCN^−^ decomposed abiotically at a slow rate, ranging from 0.058 mM to 0.02 mM per hour, proportional to the initial concentrations (Fig. 3). During the enrichment experiments, OCN^−^ concentration ranged from 0.44 to 0.72 mM over 3 to 20.5 hours. In comparison, the closest concentration for abiotic OCN^−^ decomposition experiments was 0.47 mM, with an abiotic degradation rate of 0.002 mM per hour. Over 17.5 hours (20.5 minus 3), the equivalent amount of OCN^−^ degraded would have been approximately 0.03 mM.

**Figure 3.**
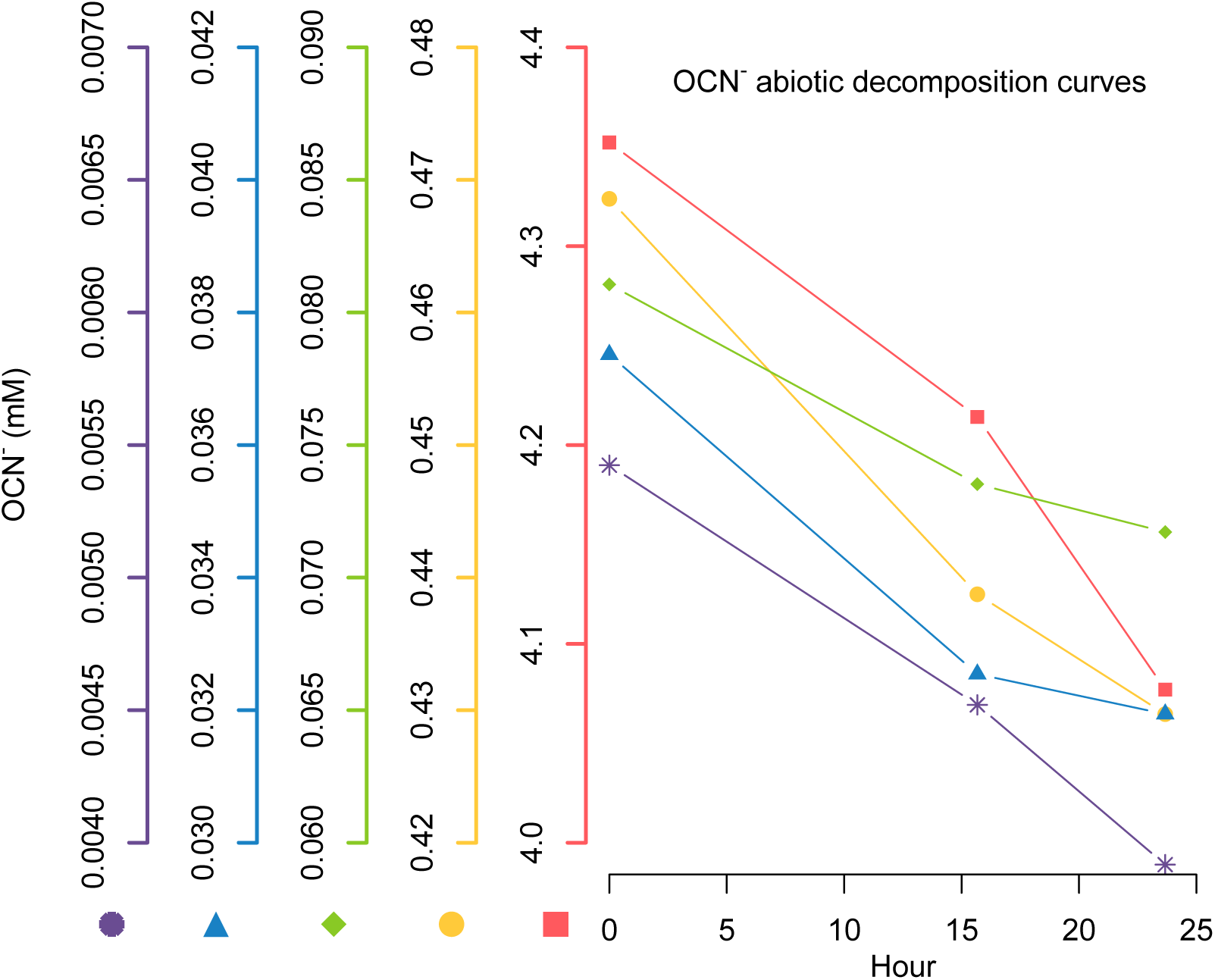
OCN^−^ concentrations during the abiotic decomposition process with initial concentrations of 4.4 (red square), 0.47 (yellow circle), 0.08 (green diamond), 0.04 (blue triangle), and 0.005 mM (purple asterisk). Higher initial OCN^−^ concentrations correspond to higher abiotic degradation rates.

### 13C– and ^15^N-labeled SCN^−^ degraders during SIP incubations

We conducted a 24-hour DNA-SIP incubation experiment to identify active SCN^−^ degraders. The extracted DNA was divided into different fractions based on their buoyant densities. The qPCR analysis of *scnC* (genes) in fractionated DNA revealed that SCN^−^ degraders via the COS pathway were successfully isotope-labelled. The maximum ratio of *scnC* in unlabeled SCN^−^ treatments was found in the fraction with a buoyant density of ∼1.70-1.72 g/mL, at both T1 and T8 time points (Figs. 4A, 4C, and 4E). In isotope-labeled S^13^C^15^N^−^ treatments, peak ratios of *scnC* were found in the fractions with buoyant densities shifting from ∼1.71 g/mL at T1 to ∼1.75-1.76 g/mL after 24 hours incubation at T8 (Figs. 4B, 4D, and 4F). Comparison of buoyant densities of *scnC* between unlabeled and labeled treatments at T1 confirms that *scnC* was not isotope-labeled at the beginning of the experiments. The buoyant densities of *scnC* between T1 and T8 in the unlabeled SCN^−^ treatments confirmed no significant C and N isotopic fractionation occurred during *scnC* synthesis. The buoyant densities of *scnC* between T1 and T8 in the isotope-labeled S^13^C^15^N^−^ treatments confirmed that *scnC* became heavier with time due to incorporation of the heavy isotopes ^13^C and ^15^N^−^ into microbial DNA.

**Figure 4.**
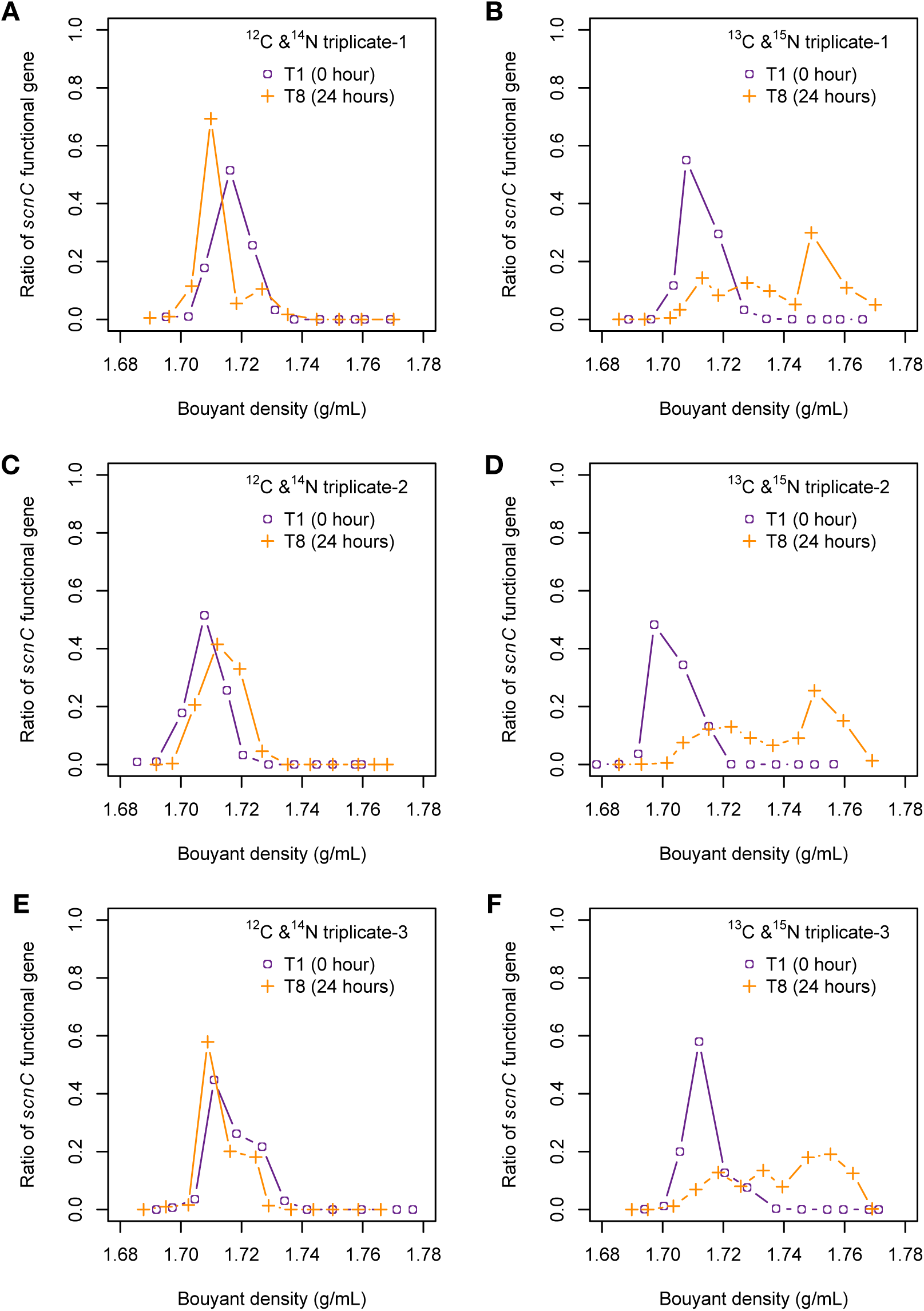
The ratios of *scnC* functional genes in the density-gradient fractionated samples recovered from DNA-SIP experiments in. (A) replicate-1 with unlabeled substrates, (B) replicate-1 with ^13^C and ^15^N labeled substrates, (C) replicate-2 with unlabeled substrates, (D) replicate-2 with ^13^C and ^15^N labeled substrates, (E) replicate-3 with unlabeled substrates, and (F) replicate-3 with ^13^C and ^15^N labeled substrates. The x-axis shows the CsCl buoyant density of the fractionated samples from lighter to heavier density. In contrast, the y-axis shows the ratios of *scnC* functional genes in all fractionated samples, quantified using qPCR. The open purple circles and orange plus symbols represent the beginning (0 hours) and end (24 hours) of the experimental period, respectively.

**Figure 5.**
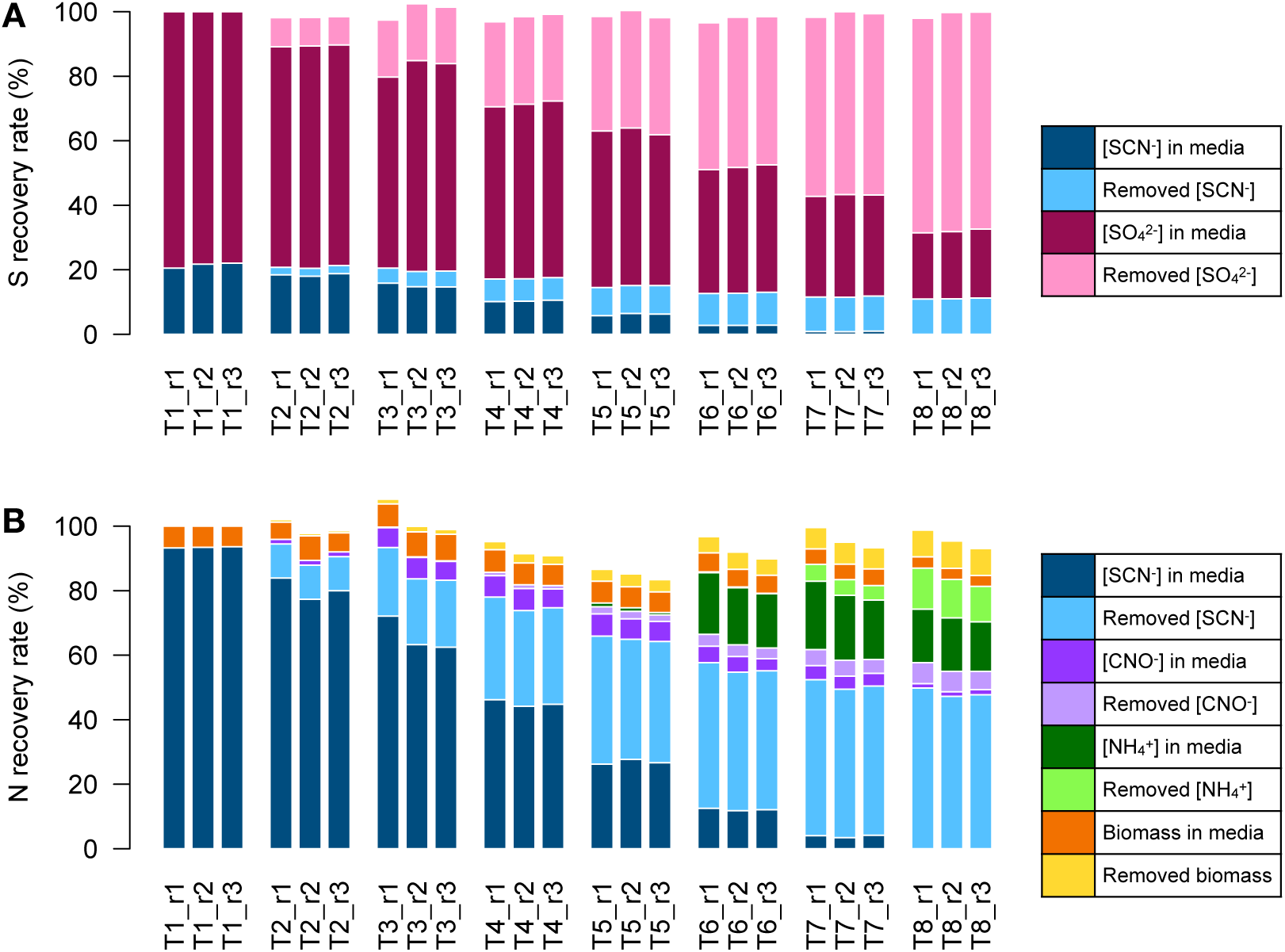
The adjusted percentage mass balance of S and N at T2 (3 hours), T3 (6 hours), T6 (17 hours), and T7 (20.5 hours) in isotope labeled. (A) replicate-1, (B) replicate-2, and (C) replicate-3 experimental groups. Compared with the N recovery (%) from the previous study, which shows in Fig. 1F only the included chemical N-compounds, the adjusted N recovery includes assimilated N in microbial biomass. The calculated and measured components are listed in **Table 2**.

### Nitrogen and sulfur mass balances

Sulfur recovery efficiency was maintained at 97.8 – 100.0% on average across the incubation period (Fig. 4A). Roughly 79% of S was comprised of SO_4_^2-^ in the media, and SCN^−^ accounted for about 21% of S-bearing compounds (there was approximately 20.3 mM SO_4_^2-^ in the media at T1). After sampling, around 11.1 % of S compounds remained as SCN^−^ at T8. Nitrogen recovery fell between 85.1% and 102.4% on average (Fig. 4B). Approximately 93% N was comprised of SCN^−^ in the media at T1, before degradation into OCN^−^ or NH_4_^+^. At the termination of experiments, N ratios for these compounds decreased to about 84%, at T8; for comparison, biomass N accounted for 7% of N compounds at T1 and 12% at T8.

## Discussion

This study presents direct and unambiguous evidence for a significant contribution (up to 70% of SCN-consumption) via the OCN^−^ pathway during SCN^−^ biodegradation by a mixed-species bioreactor microbial community. Our finding is consistent with a recent study of SCN^−^ biodegradation in a municipal wastewater treatment plant (24), based on relative quantification of mRNA reads for genes encoding the COS and OCN^−^ degradation pathways, that the OCN^−^ pathway was potentially more active (37).

Genes encoding enzymes for the COS pathway were found in our experimental microbial consortium ((23), present study). However, previously (23, 38) the OCN^−^ pathway was not observed to contribute significantly to SCN^−^ biodegradation (the maximum OCN^−^ concentration measured here was 0.75 mM, in contrast to 0.03 mM in (23). Although OCN^−^ concentrations up to ∼0.9 mM were observed previously (38), these concentrations are considered relatively low, since the starting concentration of SCN^−^ was ∼22mM (versus ∼5.5 mM in our experiment). Previous work (38) also lacked a control group to rule out the possibility of abiotic OCN^−^ generation.

Two hypotheses could explain why the OCN^−^ pathway was activated earlier than the COS pathway in our experiment: 1) physiological preference for lower temperatures and 2) the higher energy yield of the OCN^−^ pathway compared to the COS pathway. In both our previous (23, 38) and current work, the microbial consortium was stored at 4°C before the experiments. If the OCN^−^ pathway is indeed favored at lower temperatures (37), microorganisms responsible for initiating the activity of this pathway may have been pre-adapted for the onset of our experiment more than their COS pathway-utilising counterparts. This condition could favour SCN^−^ biodegradation via the OCN^−^ pathway in field-scale treatment systems where ambient or groundwater temperatures are typically below 15°C, as was the case for the SCN^−^ –contaminated tailings water inoculum for this experiment. In this study, the microbial consortium was stored and transferred to a 4°C medium prior to incubation at 30°C during the experiments. OCN⁻ was only detected after 3 hours of incubation at 30°C. However, it remains unclear whether the initial low-temperature storage provides a survival advantage to microorganisms, enabling them to degrade SCN⁻ via the OCN⁻ pathway with enhanced flexibility to cope with temperature fluctuations in the lower temperature range, which may be their optimal conditions. In contrast, the temperature of our previous experiment (23, 38) was maintained at 30°C for 16 days and 20°C for 44 days to optimize culturing conditions and promote the establishment of biofilm, before testing temperature influences on SCN^−^ degrading microbial consortia (23), supporting the hypothesis that different microbial populations utilizing different SCN^−^ –degradation pathways dominated under different temperatures. Further investigation of microbial gene expression is warranted to verify whether SCN^−^ degraders utilizing only the OCN^−^ pathway inherently prefer temperatures below 15 or 20°C.

Additionally, at the beginning of the present experiment, when the concentrations of all products were zero, the reaction quotient (Q) is also zero. In this condition, the energy yield was higher for the OCN^−^ pathway (eq 3) than the COS pathway (eqns. 1 and 2) when involving O_2_, favoring the OCN^−^ pathway. However, if O_2_ is not involved in the reaction, as suggested by a previous study (27): SCN^−^ + H_2_O → OCN^−^ + H_2_S, then the standard free energy is 194.9 kJ/mol, and the reaction will not occur spontaneously. Therefore, further investigation is required to determine whether O_2_ is involved in the OCN^−^ pathway or if microorganisms produce enzymes that can catalyze the OCN^−^ pathway.

Furthermore, microorganisms may maintain OCN^−^ concentrations, potentially leading to an underestimation of the contribution of the OCN^−^ pathway. Another previous study observed microbial control on the OCN^−^ concentration of arable soil solutions (36). In this work, urea was added into soil for OCN^−^ production, and researchers found OCN^−^ production/consumption reached equilibrium at ∼6 hours of incubation, until the end of the experiment at 30 hours (36). Here we observed OCN^−^ concentrations increased at ∼3 hours and reached a constant value at ∼13 hours until SCN^−^ was entirely degraded 24 hours. In our previous study, OCN^−^ concentrations increased at ∼50 hours and reached a constant value at ∼110 hours until SCN^−^ was degraded completely at ∼223 hours (38). Because SCN^−^ degradation can involve two pathways, the constant OCN^−^ concentrations during SCN^−^ degradation can be attributed to 1) the COS pathway replacing the OCN^−^ pathway, 2) microorganisms utilizing the OCN^−^ pathway reaching substrate saturation, or 3) an equilibrium between microbial generation and consumption of OCN^−^. We argue here that either the second or third scenario is the primary reason, with the OCN^−^ pathway continuing to contribute to SCN^−^ degradation throughout the experiments. Although we identified isotope-labelled genes encoding the COS pathway, we cannot conclude confidently that the COS pathway *replaced* the OCN^−^ pathway, since OCN^−^ may provide energy for different microorganisms (36, 39), and our previous experiments also found OCN^−^ degraders *Pseudomonas putida*, *Pseudomonas stutzeri*, and *Thiobacillus* spp. with genes encoding the COS pathway from the same field site (38). Furthermore, OCN^−^ decays naturally (Fig. 4), even though the abiotic decomposition rate is relatively slow; thus abiotic processes would decrease OCN^−^ to a detectable level during the experiments, rather than maintaining a more stable OCN^−^ concentration. Also termination of the OCN^−^ pathway activity would likely have caused OCN^−^ concentrations to begin decreasing immediately, instead of decreasing only upon complete removal of SCN^−^. Therefore, our work suggests that the OCN^−^ pathway predominates in the initial SCN^−^ biodegradation process, and continues to operate until SCN^−^ is depleted, with the COS pathway operating simultaneously rather than replacing it.

It remains to be confirmed whether the constant OCN^−^ concentration observed here was caused by microbial competition or substrate saturation. Competition among multiple OCN^−^ degraders may result in maintenance of OCN^−^ concentrations, in which the dominant metabolic guilds maintained a limited energy resource at low concentrations to inhibit others. Previous studies have observed sulfate reducers maintaining hydrogen concentrations to inhibit methanogens in anoxic sediments (40). Such maintenance only happens when energy sources are limited, which may explain why OCN^−^ concentrations were relatively similar (∼0.9 mM vs ∼0.75 mM), even if the starting concentrations of SCN^−^ in (38) was much higher than this study (∼22 mM vs ∼5.5 mM). In contrast, SCN^−^ was not a limited energy source during these experiments. Therefore, the concentrations of SCN^−^ were not maintained and OCN^−^ and COS pathways could be both present during biodegradation. On the other hand, the average SCN^−^ degradation rate in (38) was ∼0.19 mM/hr during the exponential phase. In our study, microorganisms took 24 hours to decompose ∼5.5 mM SCN^−^, which results in a ∼0.23 mM/hr degradation rate. These two rates are not much different from each other, and the contribution of the COS pathway is unknown. Therefore, we cannot rule out the possibility of substrate saturation. The difference between these two conditions, microbial competition and substrate saturation, influences the potential for OCN^−^ and SCN^−^ degradation capacity. For example, the systems have not yet reached the maximum degradation efficiency if the OCN^−^ concentration is maintained due to microbial competition. If 0.74 – 0.9 mM OCN^−^ concentrations have already reached substrate saturation levels, then this range defines the maximum bioreactor efficiency for the OCN^−^ pathway. Further investigation to evaluate the bioenergy availability in these systems and gene expression by OCN^−^ degraders would be helpful to decrypt the potential OCN^−^ degradation efficiency.

Our study experimentally confirmed the hypothesis that nitrogen from SCN-can be assimilated into microbial biomass (23). We also verified that nitrogen and carbon from SCN^−^ can be assimilated into microbial biomass via the OCN^−^ pathway, and potentially via the COS pathway. Although both ^13^C in SCN⁻ and HCO₃⁻ were provided, ^13^C incorporation into biomass occurred only through SCN⁻ degradation during the initial experimental stage. This is because the six known autotrophic carbon fixation pathways require either light (for the CBB, 3HP, and rTCA cycles) or hydrogen (for the WL, 3HP-4HB, and 4HB cycles) as energy sources (41). Our experiments were conducted in the dark, with air as the headspace, to avoid initial carbon fixation. The addition of ^13^C-labeled HCO₃⁻ was intended to identify microorganisms involved in subsequent reactions, particularly if hydrogen or sulfur were produced for the 3HP-4HB cycle. During the initial stage of the experiments, with SCN^−^ as the sole carbon and nitrogen sources, microorganisms built their biomass using carbon and nitrogen from SCN^−^. During the initial 6 hours when the OCN^−^ pathway was dominant, ^13^C and ^15^N incorporated into biomass exceeded 6 At%^15^N, indicating that SCN^−^ degraders using the OCN^−^ pathway utilized SCN^−^ derived carbon and nitrogen to build biomass.

After the initial stage, microbial cross feeding could occur, where microorganisms degrading SCN^−^ generate products such as OCN^−^ and NO_3_^−^ that serve as energy sources and biomass materials for subsequent degraders. Post 6 hours, DNA buoyant density measurements of *scnC* indicated that ^13^C and ^15^N were also incorporated into the biomass of microorganisms capable of degrading SCN^−^ via the COS pathway, with maximum ratios of *scnC* functional genes found in the fractions of CsCl gradient solutions with buoyant densities of ∼1.75 g/mL. Our results align with those of previous SIP experiments in terms of maximum ratios for unlabeled, ^15^N labeled, ^13^C labeled, and ^13^C &^15^N labeled 16S rRNA genes (DNA) having buoyant densities of ∼1.70, ∼1.71, ∼1.74, and ∼1.75 g/mL, respectively (42). This finding confirms that carbon and nitrogen were assimilated into microorganisms capable of utilizing the COS pathway. Further investigation, including measurements of COS concentrations and gene expression of the functional gene *scnC*, is required to verify the occurrence of the COS pathway in the system.

In summary, we found that microorganisms could degrade approximately 5.5 mM SCN^−^ within 24 hours using the OCN^−^ pathway, and potentially the COS pathway, initially in that order but later simultaneously until completion of the experiment. The OCN^−^ pathway contributed up to 70% of initial SCN^−^ biodegradation. Carbon and nitrogen from SCN^−^ were assimilated into the biomass of SCN^−^ degrading cells via the OCN^−^ pathway, with microorganisms capable of the COS pathway also participating in the SCN^−^ biodegrading network. Given that OCN^−^ is present mostly as an aqueous anion and COS forms a gas in most natural environments, bioreactor designs for optimizing the activity of these two enzymatic pathways for SCN-biodegradation may differ substantively. Our results offer insights both for improving bioremediation system design and understanding the relative roles of each enzymatic SCN-biodegradation pathway in biogeochemical sulfur cycling in aqueous environments.

## Material and methods

### Enrichment incubation and stable-isotope probing

Microbial inocula were collected from open-air tailings storage facilities at the Stawell Gold Mine in Victoria, Australia. The microbial consortium was enriched in a basal salt medium (pH 7.5) containing 0.5 g KSCN, 2.25 g Na_2_SO_4_, 1.00 g NaHCO_3_, 0.51 g MgSO_4_, 1.25 g CaCl_2_ · 2H_2_O, 0.1 g KCl, 1.5 g NaCl, 0.02 g NaH_2_PO_4_ · 7H_2_O, 3.5 g HEPES, and 5 mL mineral elixir solution in 1 L of medium. The mineral elixir solution (pH 8) consisted of 2.14 g Nitrilotriacetic acid (NTA), 0.1 g MnCl_2_ · 4 H_2_O, 0.3 g FeSO_4_ · 7 H_2_O, 0.2 g ZnSO_4_ · 7 H_2_O, 0.03 g CuCl_2_ · 2 H_2_O, 0.005 g AlK(SO_4_)_2_ · 12 H_2_O, 0.005 g H_3_BO_3_, 0.09 g Na_2_MnO_4_ · 2 H_2_O, 0.11 g NiSO_4_ · 6 H_2_O, 0.02 g Na_2_WO_4_ · 2 H_2_O. At late log phase of growth (OD_600_ based), the entire culture was centrifuged at 5000 g for 15 mins, and subsequently washed three times in a 4°C basal salt medium. The cells then were resuspended in 100 mL medium without KSCN and stored at 4°C for no longer than 12 hours prior to inoculation.

During the experiments, the enriched microbial consortium was separated into nine 1L serum bottles with triplicated experimental groups containing isotope-unlabeled substrate (A1, A2, and A3), triplicated experimental groups containing isotope-labeled substrates (B1, B2, and B3). Triplicated control groups (C1, C2, and C3) were sterilized at 120°C to serve as the negative controls. The isotope-labeled substrates contained KS^13^C^15^N and NaH^13^CO_3_ (Sigma, Merk, Australia) instead of KSCN and NaHCO_3_. The mixed microbial consortium was sealed with a thick butyl rubber stopper and incubated on a shaking incubator in the dark at 30°C and 120 rpm. All glassware used was acid washed using 2.5% HNO_3_ prior to use.

Homogenized samples were collected from the serum bottles at regular intervals (T1: 0 h, T2: 3 h, T3: 6 h, T4: 9 h, T5: 13 h, T6: 17 h, T7: 20.5, T8: 24 h). A total of 35 mL of liquid was removed in each sampling. Two mL of liquid sample was used for OD_600_ and SCN^−^ measurement immediately after sampling. Thirty mL of liquid samples were centrifuged, and the supernatant was passed through a 0.22 μm filter for OCN^−^, NH_4_^+^, NO_3_^−^, NO_2_^−^, and SO_4_^2-^ measurement. The pellet was either resuspended using the RNAlater Storage Solution (Sigma, Merk, Australia) and stored at –80°C for subsequent nucleic acid or dried in an 80°C oven overnight for subsequent EA-IRMS measurement.

Additional experiments to measure the OCN^−^ abiotic decomposition rates were conducted using the same basal salt medium with KOCN in concentrations of 0.005-0.2 mM and without the microbial consortium. Samples were collected at 0, 16 and 24 hours and passed through a 0.22 μm filter before determining OCN^−^ concentrations.

### Chemical parameter measurements

The growth of microorganisms was measured based on the change of OD_600_ values through a spectrophotometer (Cary 60 UV-Vis spectrophotometer, Agilent Technologies). Chemical concentrations of SCN^−^, NH_4_^+^, and NO_3_^−^ in filtered liquid samples were determined by established colorimetric protocols using a spectrophotometer (Cary 60 UV-Vis spectrophotometer, Agilent Technologies). Gravimetry via reaction with a BaCl_2_ solution was used to determine SO_4_^2-^ concentrations. The OCN^−^ concentrations were determined using an ion chromatograph (Metrohm 850 Professional IC AnCat, fitted with a Metrosep A Supp 16,150/4.0 column) with KOCN in concentrations of 0.005-0.2 mM as standards. Samples were diluted to the range of standards before measurements using a spectrophotometer or ion chromatograph. Standard deviation and the mean values were calculated and plotted.

SCN^−^ was determined by acidifying 1 mL samples or standards using 10 µL of concentrated HNO_3_, mixing with 50 µL colorimetric reagent, waiting for 30 seconds to develop color in cuvettes and measuring at 460 nm wavelength. The standards were prepared by dissolving KSCN to concentrations of 0-15 mg/L. The colorimetric reagent contained 404 g of Fe(NO_3_)_3_·9H_2_O in a 1 L solution.

For NH_4_^+^ measurements, 100 µL of filtered liquid samples or standards were mixed with 550 µL buffer solution, 400 µL salicylate solution, and 200 µL of hypochlorite solution in cuvettes. The mixed solutions were set to rest for 15 min at 37°C before homogenizing using a pipette and measuring at 650 nm. The standards were prepared by dissolving (NH_4_)_2_SO_4_ to concentrations of 0-6 mg/L. The buffer solution contained 35.8 g Na_2_HPO_4_·12H_2_O, 50 g C_4_H_4_KNaO_6_·H_2_O, and 108 g of 50% NaOH in 1 L solution. The salicylate-nitroprusside solution contained 150 g C_7_H_4_KNaO_3_ and 0.3 g Na_2_[Fe(CN)_5_NO] ·2H_2_O in a 1 L solution. The hypochlorite solution contained 6 mL of 5.25% NaOCl in 100 mL solution.

NO_3_^−^ concentrations were determined by mixing 0.025 mL of samples or standards and 0.08 mL of salicylic acid-sulfate reagent in cuvettes and waiting for 20 min to develop color, then 1.9 mL of 2 N NaOH were added and cooled to room temperature before measuring at 410 nm. The standards were prepared by dissolving KNO_3_ to concentrations of 1-300 mg/L. The salicylic acid-sulfate reagent contained 5 g of salicylic acid in 100 mL of concentrated H_2_SO_4_, made and used in the same week as the experiments and stored in the dark.

Tubes for SO_4_^2-^ measurement were dried at 80°C overnight and cooled for measuring their weights. Filtered liquid samples or standards, 3 M HCl and 1 M BaCl_2_, were mixed in the tubes with a ratio of 30:1:10 in volume. The mixed liquids were vortexed for 1 min to mix and left to precipitate overnight at room temperature. Samples then were centrifuged at 15,000 rcf for 20 minutes, and the supernatants were removed. The remained precipitates were dehydrated at 80°C overnight. The weights of precipitates were calculated by subtracting the tube weights and comparing them with the values of standard samples. The standard samples were prepared by dissolving Na_2_SO_4_ to concentrations of 2-5 g/L.

Samples for IRMS analysis were sent to the Melbourne TrACEES Platform, and the measurement was performed on a Thermo Flash 2000 HT (elemental analyser) paired to a Thermo Delta V Advantage Isotope Ratio Mass Spectrometer (IRMS). The percentage of N of the total biomass weights were calculated using TCD and calibrated against acetanilide standards (ThermoFischer, Bremen). The reference values were IVA33802174 δ^15^N_AIR_ –0.32 ‰, USGS41a δ^15^N_AIR_ 47.55 ‰, USGS43 δ^15^N_AIR_ +8.44 ‰. Data was acquired using Isodat 3.0. The δ^15^N_Air_ ‰ is defined as:

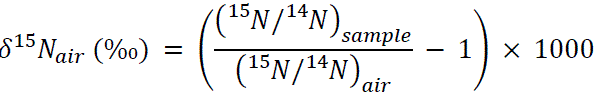

### DNA-SIP experiment set-up

DNA was extracted along with RNA following the Qiagen RNeasy PowerSoil Total RNA Isolation kit and Qiagen RNeasy PowerSoil DNA Elution Kit. All plasticware used was certified RNase-free, and the extraction environment and pipettors were cleaned using the MoBio UltraCleanTM Laboratory Cleaner and Invitrogen RNase AWAY. Roughly 2 μg of extracted DNA was added into CsCl solutions with an initial density of 1.696 g/mL. The CsCl solutions were prepared by adjusting the refractive index to 1.399 using an ATAGO-R-5000 handheld refractometer. Density gradient centrifugation was conducted using Beckman Coulter 4.9 mL OptiSeal polyallomer tubes in a Beckman Coulter VTi 90 vertical rotor at 56,200 rpm for 24 hours at 20°C. Centrifuged gradients were fractionated into 25∼200 μL volumes using a Beckman Coulter fraction recovery system and CBS-Scientific MPP-100 Mini Peristaltic Pump. The buoyant density of each collected fraction was measured using 25 μL aliquots and a refractometer to determine the refractive indexes. DNA was precipitated from CsCl solution overnight by adding two volumes (400 μL in this case) of PEG 6000 in 1.6 M NaCl solutions. The precipitated DNA was then centrifuged at 13,000 rpm at 4°C for 20 minutes and washed using 70% ethanol before being dissolved in 30 μL of sterile water.

### qPCR gene quantification

The qPCR (quantitative polymerase chain reaction) for *scnC* functional genes quantification of DNA-SIP gradients was performed using Bio-Rad SsoAdvanced Universal SYBR Green Supermix with primer set *scn*-1 (5’-GTN GCN MRN GCN TGG BTN GAY CC-3’) and *scn*-2 (5’-GGI CKI WSI GGI ADI CAN ADR TA-3’) (41) with annealing temperature 56°C.

### Data processsing

The standard Gibbs free energy values were calculated using the R software (43) and the package CHNOSZ v.2.1.0 (44), with the OBIGT thermodynamic database. The standard free energy value of COS was manually input using data from a published reference (45).

The ratios of *scnC* functional genes in each collected SIP density gradient fraction are expressed as the number of calculated *scnC* gene copies of the certain fraction over the sum of *scnC* gene copies in all the fractions from the same CsCl gradient tube.

To investigate S and N recovery rates, we calculated all components in moles. The S components include SCN^−^ and SO_4_^2-^ in current and removed media. The media contained SO_4_^2-^ in different chemical forms, including 15.8 mM Na_2_SO_4_, 4.2 mM Mg_2_SO_4_, 0.005 mM FeSO_4_ · 7H_2_O, 0.0001 mM KAl(SO_4_)_2_ · 12H_2_O, 0.002 mM NiSO_4_ · 6H_2_O, and 0.003 mM ZnSO_4_ · 7H_2_O. Therefore, SO_4_^2-^ comprised the dominant S compound since the beginning of the incubation experiments. The N compounds included SCN^−^, OCN ^−^, NH_4_^+^, and biomass in current and removed media. The biomass of N was calculated based on weight, and then the N% was measured using IRMS for samples at T2, T3, T6, and T7. The biomass of N at T1, T4, T5, and T8 was calculated by extrapolation and interpolation using the trendline based on the measured biomass N. The molar values of removed chemicals and the weight of removed biomass were calculated based on the concentrations measured at the previous time points. For example, the SCN^−^ components at T3 include SCN^−^ in T3 media and 35 mL media removed at T1 and 35 mL at T2. The N and S recovery rates were calculated based on the total N and S moles at a particular timepoint compared to T1 values. The contribution of the OCN⁻ pathway is expressed as the proportion of SCN⁻ converted to OCN⁻, calculated by dividing the OCN⁻ concentration by the SCN^−^ concentration.

## Acknowledgements

The authors would like to acknowledge David Coe, Will Wettenhall, Troy Cole, Peter Wemyss, and Cameron Hope (Stawell Gold Mine, Australia) for their support of this project and access to mine tailings storage facilities. We also thank Jay Black (The University of Melbourne, Australia) for his help with ion chromatography; Farhad Shafiei and Lukas Pajank (The University of Melbourne, Australia) for their help with spectrophotometry; Prof. Jim He, Dr. Hang-Wei Hu, Qing Xie, and Chaoyu Li (The University of Melbourne) for their invaluable advice with stable isotope probing and help with ultracentrifugation; and Michael Hall (Melbourne TrACEES Platform, The University of Melbourne) for his help with IRMS measurements. This research was funded by Australian Research Council Linkage Project grant #LP160100866.

## References

1. Gupta N, Balomajumder C, Agarwal V. 2010. Enzymatic mechanism and biochemistry for cyanide degradation: a review. 1–3. Journal of hazardous materials 176:1–13.

2. Jeffrey MI, Breuer PL. 2000. The cyanide leaching of gold in solutions containing sulfide. Minerals Engineering 13:1097–1106.

3. Johnson CA. 2015. The fate of cyanide in leach wastes at gold mines: An environmental perspective. Applied geochemistry 57:194–205.

4. Zhang C, Wang X, Jiang S, Zhou M, Li F, Bi X, Xie S, Liu J. 2021. Heavy metal pollution caused by cyanide gold leaching: a case study of gold tailings in central China. Environmental Science and Pollution Research 28:29231–29240.

5. Staib C, Lant P. 2007. Thiocyanate degradation during activated sludge treatment of coke-ovens wastewater. Biochemical engineering journal 34:122– 130.

6. Granda M, Blanco C, Alvarez P, Patrick JW, Menendez R. 2014. Chemicals from coal coking. Chemical Reviews 114:1608–1636.

7. Li C, Li G, Zhang S, Wang H, Wang Y, Zhang Y. 2018. Study on the pyrolysis treatment of HPF desulfurization wastewater using high-temperature waste heat from the raw gas from a coke oven riser. RSC advances 8:30652–30660.

8. Li L, Yue F, Li Y, Yang A, Li J, Lv Y, Zhong X. 2020. Degradation pathway and microbial mechanism of high-concentration thiocyanate in gold mine tailings wastewater. RSC Advances 10:25679–25684.

9. Cho Y, Cattrall RW, Kolev SD. 2018. A novel polymer inclusion membrane based method for continuous clean-up of thiocyanate from gold mine tailings water. Journal of hazardous materials 341:297–303.

10. Heming TA, Thurston RV, Meyn EL, Zajdel RK. 1985. Acute toxicity of thiocyanate to trout. Transactions of the American Fisheries Society 114:895– 905.

11. Lanno RP, Dixon DG. 1994. Chronic toxicity of waterborne thiocyanate to the fathead minnow (pimephales promelas): A partial life-cycle study. Environmental Toxicology and Chemistry: An International Journal 13:1423– 1432.

12. Boening DW, Chew CM. 1999. A critical review: general toxicity and environmental fate of three aqueous cyanide ions and associated ligands. Water, air, and soil pollution 109:67–79.

13. Brown P, Morra M. 1993. Fate of ionic thiocyanate (SCN-) in soil. Journal of agricultural and food chemistry 41:978–982.

14. Kurashova I, Halevy I, Kamyshny Jr A. 2018. Kinetics of decomposition of thiocyanate in natural aquatic systems. Environmental science & technology 52:1234–1243.

15. Spurr LP, Watts MP, Gan HM, Moreau JW. 2019. Biodegradation of thiocyanate by a native groundwater microbial consortium. PeerJ 7:e6498–e6498.

16. Huddy RJ, Kantor R, Van Zyl W, Van Hille RP, Banfield JF, Harrison STL. 2015. Analysis of the microbial community associated with a bioprocess system for bioremediation of thiocyanate-and cyanide-laden mine water effluents, p. 614–617. *In*. Trans Tech Publ.

17. Watts MP, Moreau JW. 2018. Thiocyanate biodegradation: harnessing microbial metabolism for mine remediation. Microbiology Australia 39:157–161.

18. Laurberg P, Pedersen IB, Carlé A, Andersen S, Knudsen N, Karmisholt J. 2009. The relationship between thiocyanate and iodine. Comprehensive handbook of iodine: nutritional, biochemical, pathological and therapeutic aspects Amsterdam: Elsevier 275–281.

19. U.S. EPA. 2012. Provisional Peer-Reviewed Toxicity Values for Thiocyanic Acid. Washington.

20. Watts MP, Gan HM, Peng LY, Lê Cao K-A, Moreau JW. 2017. In situ stimulation of thiocyanate biodegradation through phosphate amendment in gold mine tailings water. Environmental science & technology 51:13353–13362.

21. Huddy RJ, Van Zyl AW, Van Hille RP, Harrison ST. 2015. Characterisation of the complex microbial community associated with the ASTER^TM^ thiocyanate biodegradation system. Minerals Engineering 76:65–71.

22. Shafiei F, Watts MP, Pajank L, Moreau JW. 2021. The effect of heavy metals on thiocyanate biodegradation by an autotrophic microbial consortium enriched from mine tailings. Applied Microbiology and Biotechnology 105:417–427.

23. Watts MP, Spurr LP, Lê Cao K-A, Wick R, Banfield JF, Moreau JW. 2019. Genome-resolved metagenomics of an autotrophic thiocyanate-remediating microbial bioreactor consortium. Water research 158:106–117.

24. Broman E, Jawad A, Wu X, Christel S, Ni G, Lopez-Fernandez M, Sundkvist J- E, Dopson M. 2017. Low temperature, autotrophic microbial denitrification using thiosulfate or thiocyanate as electron donor. Biodegradation 28:287–301.

25. Zhao F, Zhang Q, He L, Yang W, Si M, Liao Q, Yang Z. 2023. Molecular level insight of thiocyanate degradation by Pseudomonas putida TDB-1 under a high arsenic and alkaline condition. Science of The Total Environment 874:162578.

26. Mekuto L, Ntwampe SKO, Kena M, Golela MT, Amodu OS. 2016. Free cyanide and thiocyanate biodegradation by Pseudomonas aeruginosa STK 03 capable of heterotrophic nitrification under alkaline conditions. 1. 3 Biotech 6:6–6.

27. Watts MP, Moreau JW. 2016. New insights into the genetic and metabolic diversity of thiocyanate-degrading microbial consortia. Applied microbiology and biotechnology 100:1101–1108.

28. Li L, Lollar BS, Li H, Wortmann UG, Lacrampe-Couloume G. 2012. Ammonium stability and nitrogen isotope fractionations for NH4+–NH3 (aq)– NH3 (gas) systems at 20–70 C and pH of 2–13: Applications to habitability and nitrogen cycling in low-temperature hydrothermal systems. Geochimica et Cosmochimica Acta 84:280–296.

29. Ryu B-G, Kim W, Nam K, Kim S, Lee B, Park MS, Yang J-W. 2015. A comprehensive study on algal–bacterial communities shift during thiocyanate degradation in a microalga-mediated process. Bioresource Technology 191:496– 504.

30. Kantor RS, Huddy RJ, Iyer R, Thomas BC, Brown CT, Anantharaman K, Tringe S, Hettich RL, Harrison STL, Banfield JF. 2017. Genome-resolved meta-omics ties microbial dynamics to process performance in biotechnology for thiocyanate degradation. 5. Environmental science & technology 51:2944–2953.

31. Svoronos PDN, Bruno TJ. 2002. Carbonyl Sulfide: A Review of Its Chemistry and Properties. Ind Eng Chem Res 41:5321–5336.

32. Aydin M, Britten GL, Montzka SA, Buizert C, Primeau F, Petrenko V, Battle MB, Nicewonger MR, Patterson J, Hmiel B, Saltzman ES. 2020. Anthropogenic Impacts on Atmospheric Carbonyl Sulfide Since the 19th Century Inferred From Polar Firn Air and Ice Core Measurements. JGR Atmospheres 125:e2020JD033074.

33. Yamasaki M, Matsushita Y, Namura M, Nyunoya H, Katayama Y. 2002. Genetic and immunochemical characterization of thiocyanate-degrading bacteria in lake water. Applied and environmental microbiology 68:942–946.

34. Berben T, Overmars L, Sorokin DY, Muyzer G. 2017. Comparative genome analysis of three thiocyanate oxidizing Thioalkalivibrio species isolated from soda lakes. Frontiers in microbiology 8:254–254.

35. Davidson D. 2018. Development of a specific PCR to detect thiocyanate-oxidizing Thioalkalivibrio strains in their environment. Amsterdam.

36. Mooshammer M, Wanek W, Jones SH, Richter A, Wagner M. 2021. Cyanate is a low abundance but actively cycled nitrogen compound in soil. 1. Communications Earth & Environment 2:1–10.

37. Ni G, Canizales S, Broman E, Simone D, Palwai VR, Lundin D, Lopez-Fernandez M, Sleutels T, Dopson M. 2018. Microbial community and metabolic activity in thiocyanate degrading low temperature microbial fuel cells. Frontiers in microbiology 9:2308–2308.

38. Watts MP, Spurr LP, Gan HM, Moreau JW. 2017. Characterization of an autotrophic bioreactor microbial consortium degrading thiocyanate. 14. Applied microbiology and biotechnology 101:5889–5901.

39. Palatinszky M, Herbold C, Jehmlich N, Pogoda M, Han P, von Bergen M, Lagkouvardos I, Karst SM, Galushko A, Koch H. 2015. Cyanate as an energy source for nitrifiers. 7563. Nature 524:105–108.

40. Hoehler TM, Alperin MJ, Albert DB, Martens CS. 1998. Thermodynamic control on hydrogen concentrations in anoxic sediments. 10. Geochimica et cosmochimica acta 62:1745–1756.

41. Zhao T, Li Y, Zhang Y. 2021. Biological carbon fixation: a thermodynamic perspective. Green Chemistry 23:7852–7864.

42. Cupples AM, Shaffer EA, Chee-Sanford JC, Sims GK. 2007. DNA buoyant density shifts during 15N-DNA stable isotope probing. 4. Microbiological research 162:328–334.

43. R Core Team R. 2013. R: A language and environment for statistical computing.

44. Dick JM. 2019. CHNOSZ: Thermodynamic calculations and diagrams for geochemistry. Frontiers in Earth Science 7:180.

45. Haynes WM. 2016. CRC handbook of chemistry and physics. CRC press. https://www.taylorfrancis.com/books/mono/10.1201/9781315380476/crc-handbook-chemistry-physics-william-haynes. Retrieved 16 July 2024.

